# Anamnestic Humoral Correlates of Immunity Across SARS-CoV-2 Variants of Concern

**DOI:** 10.1101/2022.06.19.496718

**Authors:** Ryan P. McNamara, Jenny S. Maron, Harry L. Bertera, Julie Boucau, Vicky Roy, Amy K. Barczak, The Positives Study Staff, Nicholas Franko, Jonathan Z. Li, Jason S. McLellan, Mark J. Siedner, Jacob E. Lemieux, Helen Y. Chu, Galit Alter

**Affiliations:** Ragon Institute of MGH, MIT, and Harvard, Cambridge, MA, USA; Department of Medicine, Massachusetts General Hospital, Boston, MA, USA; Harvard Medical School, Boston, MA, USA; Division of Allergy and Infectious Diseases, University of Washington, Seattle, WA, USA; Department of Medicine, Brigham and Women’s Hospital; Department of Molecular Biosciences, University of Texas at Austin, Austin, TX, USA; The Broad Institute, Cambridge, MA, USA

**Keywords:** Vaccines, COVID-19, breakthrough, SARS-CoV-2, Omicron, Delta, antibodies

## Abstract

While immune correlates against SARS-CoV-2 are typically defined at peak immunogenicity following vaccination, immunologic responses that expand selectively during the anamnestic response following infection can provide mechanistic and detailed insights into the immune mechanisms of protection. Moreover, whether anamnestic correlates are conserved across VoCs, including the Delta and more distant Omicron variant of concern (VoC), remains unclear. To define the anamnestic correlates of immunity, across VOCs, we deeply profiled the humoral immune response in individuals recently infected with either the Delta or Omicron VoC. While limited acute N-terminal domain and RBD-specific immune expansion was observed following breakthrough, a significant immunodominant expansion of opsinophagocytic Spike-specific antibody responses focused largely on the conserved S2-domain of SARS-CoV-2 was observed 1 week after breakthrough infection. This S2-specific functional humoral response continued to evolve over 2-3 weeks following both Delta and Omicron breakthrough infection, targeting multiple VoCs and common coronaviruses. These responses were focused largely on the fusion peptide 2 and heptad repeat 1, both associated with enhanced rates of viral clearance. Taken together, our results point to a critical role of highly conserved, functional S2-specific responses in the control of SARS-CoV-2 infection, across VOCs, and thus humoral response linked to virus attenuation can guide next-generation generation vaccine boosting approaches to confer broad protection against future SARS-CoV-2 VoCs.

## Introduction

Despite the remarkable vaccine efficacy observed in phase 3 SARS-CoV-2 vaccine trials, the waning of vaccine-conferred immunity and the emergence of neutralizing antibody-resistant variants of concern (VoCs), such as the Delta (B.1.612) and Omicron (B.1.529), led to a rapid increase in transmission events globally (*1-4*). Yet, severe disease and death did not increase concomitantly suggesting that additional post-transmission blocking immune responses contribute to control and clearance of infection once it has occurred (*5*). However, the precise immunologic correlates of immunity, following breakthrough infections remain incompletely defined. Moreover, whether these correlates differ across VOCs, that exhibit striking differences in sequence, is unclear.

Neutralizing antibody responses were tightly linked to protective immunity in early mRNA vaccine phase 3 trials, at a time when the dominant circulating strain was largely matched to the vaccine antigen-insert sequence (*6-8*). However, with the introduction of more neutralization-resistant VoCs, the predictive power of neutralization diminished (*2*). Despite VoC evasion of neutralization (*9-11*), both T-cells (*12*) and binding antibodies (*13*) were proposed as alternate immune mechanisms that could mediate post-treatment control and clearance of infection. Post-challenge correlates analyses in non-human primate vaccine models pointed to a rapid humoral anamnestic response, linked to the rapid expansion of antibody-secreting cells within the respiratory tract, giving rise to robust renewed pools of antibodies that may contribute to control and clearance of infection (*14*). Yet, whether these antibodies contribute to attenuation of disease via the simple neutralization and blockade of further spread or via the recruitment of the antiviral activity of the local immune system, via Fc-effector functions, is unknown. Moreover, whether the specificity and functional activity of the anamnestic response that evolves following VoC infections are conserved may point to common or distinct mechanisms of attenuation of disease.

Thus, while vaccine-induced immune correlates of protection are most often focused on the identification of immunologic responses at peak-immunogenicity, these immunologic markers may have limited consequence as mechanistic correlates of immunity, as these responses may wane at the time of environmental exposure. Conversely, immunologic signatures following exposure may provide critical insights into the anamnestic response that is key to control and clearance of the infection. Thus, here we deeply profiled the humoral immune response in a cohort of individuals with a recent, documented Delta or Omicron SARS-CoV-2 infection. Systems Serology profiling revealed a rapid expansion of Fc-receptor binding and opsinophagocytic humoral immune responses across the VoC breakthroughs with a consistent preference for expansion for the S2 subdomain of Spike, focused on the fusion peptide 2 and heptad repeat 1, that tracked with enhanced viral clearance. These data point to a critical role for S2-specific immunity as a key correlate of immunity across VoCs breakthrough infections.

## Results

### Breakthrough COVID-19 Elicits Spike Sub-domain Humoral Responses

The perpetual emergence of new SARS-CoV-2 variants of concern have led to repeated waves of viral breakthrough infections, even in recently vaccinated individuals (*5*). However, vaccine-induced immunity continues to provide protection against severe disease and death, as evidenced by the severe disease caused by both Delta and Omicron preferentially in unvaccinated populations (*4, 15-17*). Emerging data point to the potential importance of the anamnestic response as a key contributor to the resolution of infection (*18*). Given the association between non-neutralizing antibody effector profiles and natural resolution of severe disease (*19*), we performed systems serology on sera from individuals who had completed their vaccine series and had subsequent documented Delta (n = 37) or Omicron (n = 23) VoC breakthrough, both 1 week and 2-3 weeks post-infection (means = 6.3 ± 2.8, and 18.8 ± 2.9 days, respectively) aimed at defining the specific humoral properties associated with the resolution of infection. All individuals had received the primary 2 dose series of an mRNA vaccine (Pfizer/BNT162b2 = 24, and Moderna mRNA-1273 = 16). Delta and Omicron breakthroughs ranged from 5 – 357 days from vaccination and 0 – 12 days from symptom onset. Given the significant antigenic distance between Delta and Omicron within the Spike protein, we also sought to define whether anamnestic correlates were consistent across VoCs (*20*).

The rapid expansion of Spike-specific, receptor-binding domain (RBD) and N-terminal domain (NTD) specific immune responses have been proposed as potential acute anamnestic correlates of immunity following vaccine breakthrough. However, no expansion was observed in IgG responses to the full-length Spike-specific, NTD, or RBD within the first week following breakthrough infection. Moreover, *de novo* IgM, IgG3, and IgA responses to full Spike were also not observed (**Figure 1A-C**). Conversely, IgM responses expanded to the conserved S2 domain of the Spike antigen within the first week following both Delta and Omicron breakthroughs. Interestingly, S2-specific IgA also expanded following Delta (**Figure 1D**), but not Omicron breakthrough. These data point to an unexpected, and selective expansion of *de novo* humoral immune responses to the highly conserved S2 domain of the Spike antigen as a key correlate of immunity following vaccine breakthrough infection.

**Figure 1.**
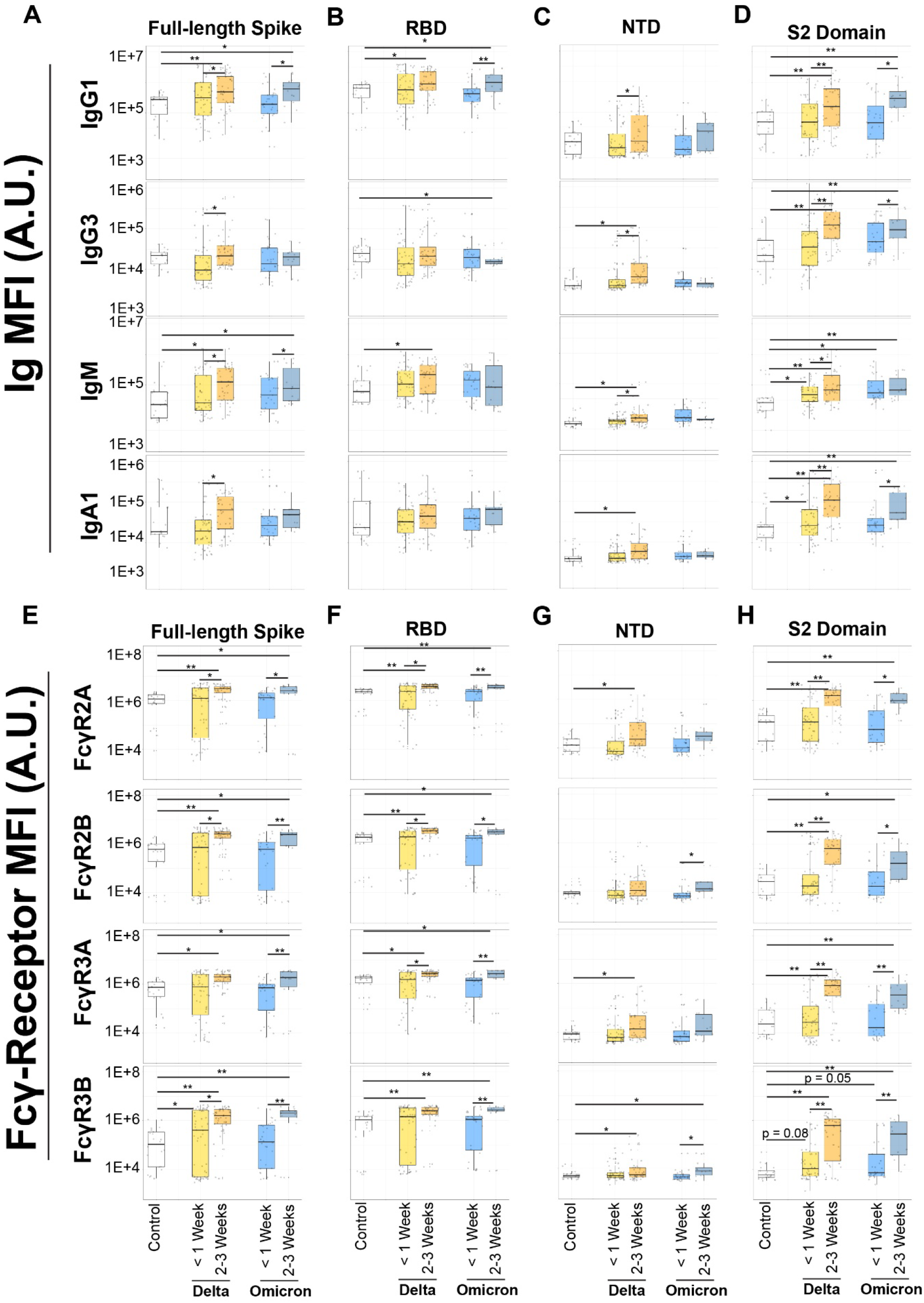
Recognition and expansion of Spike and Spike subdomains post-breakthrough. (A) Full-length Spike (D614G) was assayed for antibody recognition in non-breakthrough, vaccinated controls (white, column 1), vaccinated Delta <1 Week (yellow, column 2) or 2-3 Weeks post-breakthrough (orange, column 3), and vaccinated Omicron <1 week (blue, column 4) or 2-3 Weeks post-breakthrough (dark blue, column 5). (B) Same as A, but for the receptor-binding domain (RBD). (C) Same as B, but for the N-terminal domain (NTD). (D) Same as B, but for the S2 domain. (E) Same as A, but for Fcγ-receptor (FcγR) recognition of full-length Spike. (F) Same as E, but for the RBD. (G) Same as F, but for the NTD. (H) Same as F, but for the S2 domain. * = p < 0.05, and ** = p < 0.01 for all panels.

After 2-3 weeks, a broader expansion was observed in the full Spike-specific IgG1 response compared to uninfected vaccinees in both Delta and Omicron breakthrough infection. Spike-specific IgG3 and IgA responses increased compared to the initial post-breakthrough timepoint (**Figure 1A**) but were not higher than levels observed in controls. A similar IgG1 RBD-specific expansion was observed across both Delta and Omicron cases after 2-3 weeks of infection (**Figure 1B**). Limited expansion was observed in NTD across isotypes and was exclusively observed in Delta breakthroughs (**Figure 1C**), likely associated with the greater conservation in NTD across the vaccine insert and the Delta NTD sequence. Conversely, 2-3 weeks post-breakthrough, a highly significant expansion was observed in the S2-specific response in IgG1, IgG3, IgM, and IgA1, over time as well as compared to uninfected controls (**Figure 1D**). These data point to a highly selective and preferential continued anamnestic maturation of the S2-specific response following both Delta and Omicron breakthrough infections in vaccinees.

Binding antibodies alone do not mediate immunologic clearance, thus we next probed whether the anamnestic expansion of antibodies also possessed the ability to bind to Fc-receptors (FcR), key to leveraging non-neutralizing innate immune effector functions (*21, 22*). Interestingly, after 7 days of infection, FcR binding antibodies did not emerge to most regions of the Spike antigen. However, a significant Spike-specific FcγR3B binding response was detected in Delta breakthroughs (**Figure 1E**), and a trend towards significance was noted to the S2-domain in both Delta and Omicron breakthrough cases (**Figure 1H**).

However, 2-3 weeks following breakthrough, Spike-specific antibodies able to bind to FcRs expanded in both individuals that experienced a Delta and Omicron breakthrough (**Figure 1E**). An expansion of activating, opsinophagocytic FcγR2A and inhibitory FcγR2B binding RBD-specific antibodies was observed in both Delta and Omicron breakthroughs (**Figure 1F**). A more limited expansion of inhibitory FcγR2B and neutrophil-specific FcγR3B binding NTD-specific antibodies was observed (**Figure 1G**). Conversely, S2-specific FcR binding expanded highly significantly across all FcRs (**Figure 1H**). Collectively, these data point to an expansion of FcR binding to several Spike subdomains, including a selective expansion across isotype, subclass, and FcR binding to the highly conserved S2-domain of the Spike antigen (Supplementary Figure 1).

### Recognition of S2 is Expanded in Breakthrough Cases

To next directly compare the extent of the anamnestic expansion across Spike domains across the breakthrough cases, we examined the fold increase in IgG1 levels across subdomains of Spike, VoCs, and common human coronaviruses (HCoVs). While humoral immune responses increased to all domains in Delta breakthroughs a clear and more significant expansion was observed in S2-specific IgG1 titers (**Figure 2A**). A similar highly significant expansion was observed in S2-specific IgG1 responses in Omicron breakthroughs; however a more limited expansion was observed in RBD and NTD-specific responses, likely due to greater sequence disparity between the original vaccine antigen and Omicron within these domains (**Figure 2B**). These data point to S2 as the immunodominant anamnestic target of the humoral immune response following Delta and Omicron breakthrough.

**Figure 2.**
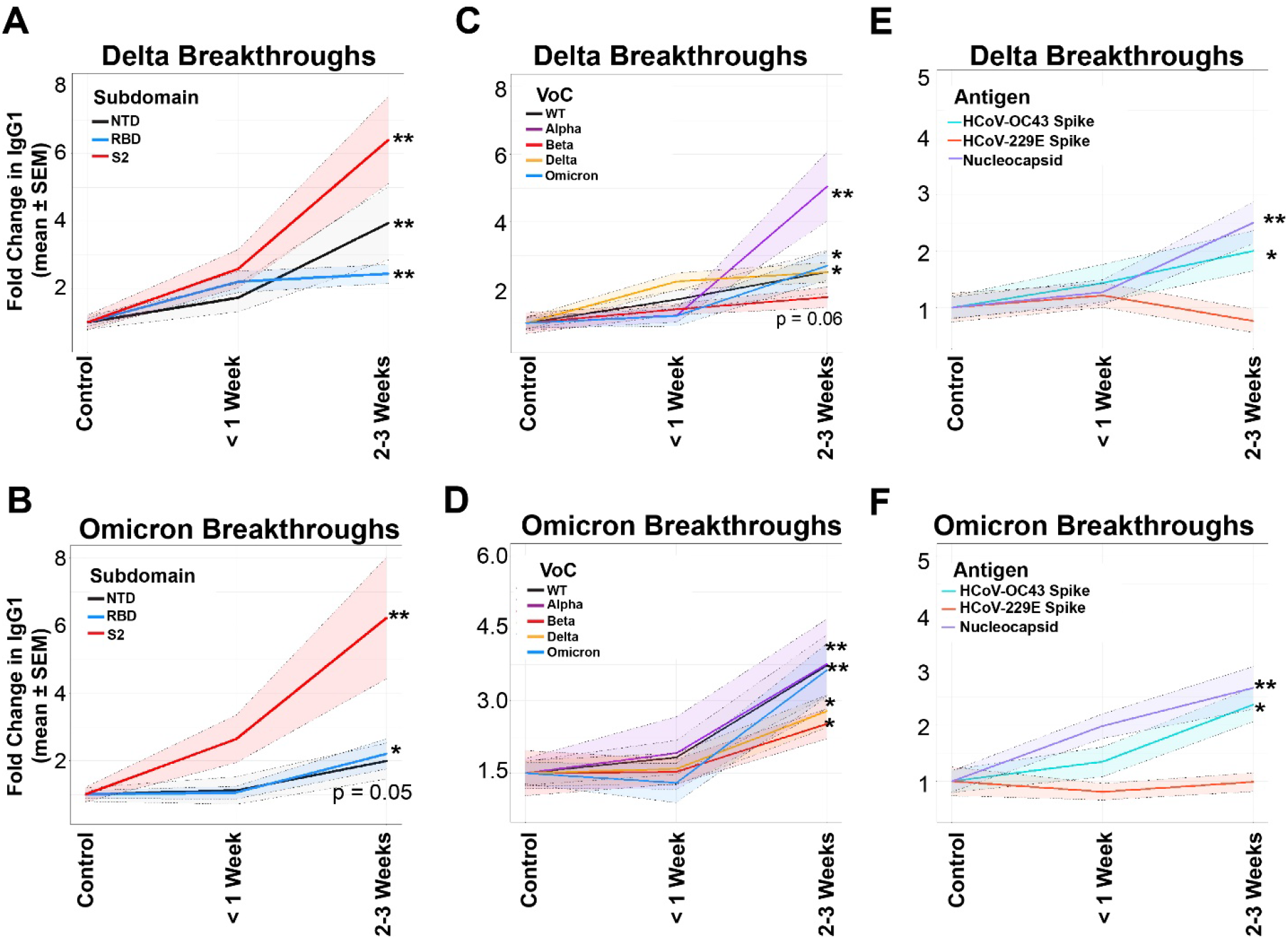
Conserved regions of Spike are selectively expanded in breakthrough cases. (A) Fold changes in IgG1 binding of the subdomains of Spike (NTD in black, RBD in blue, and S2 in red) for vaccinated, Delta breakthrough cases at <1 week or 2-3 Weeks post-breakthrough. (B) Same as A, but for vaccinated Omicron breakthrough cases. (C) Fold changes in IgG1 binding of the full-length Spikes from VoC in Delta breakthrough cases. (D) Same as C, but for Omicron breakthrough cases. (E) Fold changes in IgG1 binding of the Spike of common CoV Spikes in Delta breakthrough infections at < 1 Week or 2-3 Weeks post-breakthrough. SARS-CoV-2 nucleocapsid (N) is used as a control for infection. (F) Same as C, but for Omicron breakthroughs. * = p < 0.05, and ** = p < 0.01 for all panels.

Comparison of the breadth of the fold anamnestic expansion across Spike VoCs pointed to an overall similar expansion of response to all VoCs in Delta breakthrough cases (**Figure 2C**), with the exception of the Alpha variant Spike-specific response that expanded preferentially. Conversely, all VoC Spike-specific responses expanded significantly in Omicron breakthrough infection (**Figure 2D**), likely due to the highly divergent nature of the Omicron spike that drove enhanced B cell recruitment and affinity maturation. Moreover, analysis of responses to two common human coronaviruses (HCoVs), α (229E) and β (OC-43), revealed a selective expansion of cross-β-CoV immunity in Delta and Omicron breakthrough cases, likely due to the closer phylogenetic relation of β-CoV to SARS-CoV-2, particularly in the S2-domain (*23*) (**Figure 2E-F**). Moreover, analysis of the overall humoral coordination within the SARS-CoV-2 responses revealed similar expansion profiles across Delta and Omicron breakthroughs (Supplementary Figure 2). Thus, overall, breakthroughs of Delta or Omicron VoCs are associated with similar anamnestic immunity, marked by a novel expansion of cross-VoC and β-CoV immunity, likely directed at the conserved S2 region of SARS-CoV-2.

### S2 is the Subdomain that Drives Humoral Expansion Post-breakthrough

To next define a minimal multivariate signature of Delta or Omicron breakthrough infection among vaccinated individuals, we performed partial least squares discriminant analysis (PLS-DA) on antibody responses collected in breakthrough cases 2-3 weeks following infection. Significant heterogeneity was observed in multivariate antibody profiles across Delta (**Figure 3A**) and Omicron (**Figure 3B**) breakthrough cases compared to uninfected vaccinated controls. In fact, vaccinated controls demonstrated a highly homogeneous antibody profile, with controls consistently clustering in a small area of the multivariate space in both comparisons (**Figure 3A-B**). Conversely, both Delta breakthrough (**Figure 3A**) and Omicron breakthrough (**Figure 3B**) segregated nearly completely from uninfected control profiles based on Fc-profiling data. To gain further insights into the specific features that were most distinct across breakthrough and uninfected vaccine profiles, a variable importance plot was generated, highlighting the minimal features that were required to resolve antibody profiles (**Figure 3C**). Strikingly, of the NTD-specific antibody responses, only NTD-specific IgA responses were preferentially enriched among both Delta and Omicron breakthroughs. Conversely, distinct RBD-specific FcR binding responses, but not isotype titers, were highly discriminatory of breakthrough cases compared to vaccinated controls, marked by higher RBD-specific FcγR3B and FcγR2A in Omicron breakthroughs and FcγR3A and FcγR2B responses selectively expanded in Delta breakthroughs.

**Figure 3.**
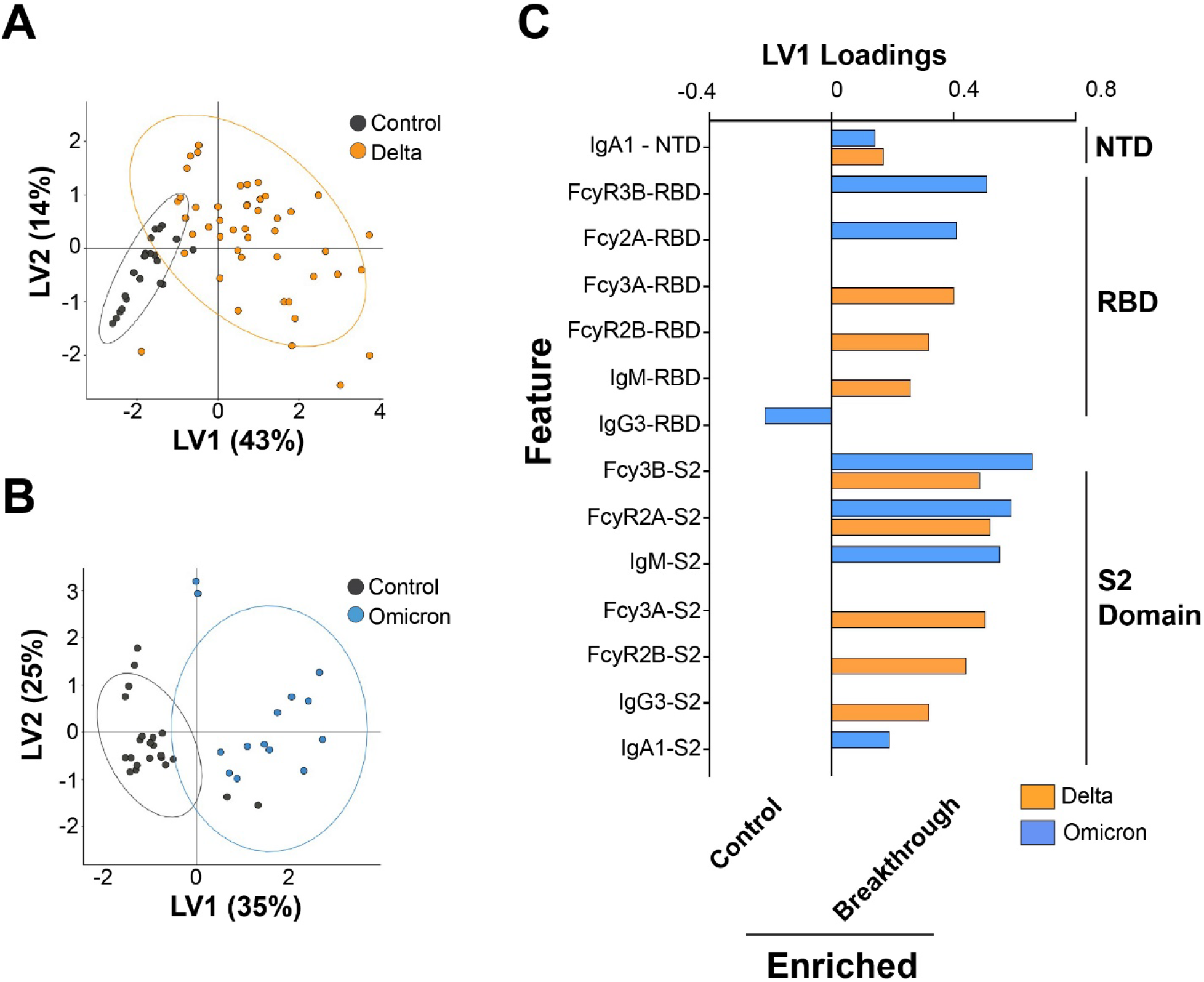
Expansion of S2 recognition is a marker for breakthrough COVID-19. (A) Partial least squares determinant analysis (PLS-DA) clustering of vaccinated, controls (black), and vaccinated Delta breakthrough (orange) immunological signatures. Shown are the clusters at 2-3 Weeks post-breakthrough. (B) Same as A, but for vaccinated Omicron breakthrough (blue) immunological signatures. (C) Ranked sum of identified features in the PLS-DA by subdomain. Shown is the distance on LV1, which was the major axis of separation by both Delta and Omicron breakthroughs at 2-3 Weeks post-breakthrough.

Instead, both S2-specific isotype titers and FcR binding antibodies were preferentially expanded across Delta and Omicron breakthrough cases compared to uninfected vaccinees. Specifically, a coordinated expansion of neutrophil-specific FcγR3B and opsinophagocytic FcγR2A binding antibodies was selectively expanded in both Delta and Omicron breakthrough profiles. Additionally, S2-specific IgM and IgA responses expanded preferentially in Omicron breakthroughs, and IgG3, FcγR3A, and FcγR2B levels were selectively observed in Delta breakthrough cases. Moreover, a near-identical pattern of S2 expansion was observed in a second, independent cohort across Delta and Omicron breakthroughs after 14 days (Supplementary Figure 3). These data collectively, highlight the broader selective expansion of functional FcR binding responses, largely focused on S2, across both Delta and Omicron breakthrough infections.

### Breakthrough infection drives a functional S2-specific humoral immune expansion

Whether the expansion of S2-specific immunity is simply a biomarker of exposure to the virus or a mechanistic correlate of immunity is unclear. However, binding alone is insufficient to drive antiviral control or clearance, thus we next aimed to determine the functional capacity of the breakthrough anamnestic response. Critically, among the antibody effector functions, we previously observed a preferential expansion of Spike-specific opsinophagocytic activity in natural survivors of severe disease (*22*) and thus we profiled the Spike- and S2-specific antibody-dependent monocyte and neutrophil phagocytic profile. Limited expansion of Spike-specific antibody-dependent cellular monocyte phagocytic (ADCP) activity was observed across the Delta and Omicron breakthrough cases (**Figure 4A**). Conversely, we observed a limited S2-specific ADCP expansion within the first week of breakthrough infection in Delta breakthroughs, but a highly significant expansion of S2-specific ADCP 2-3 weeks following breakthrough. Conversely, S2-specific ADCP was already significantly expanded in Omicron-infections by day 7 and continued to expand over the next few weeks following breakthrough infection.

**Figure 4.**
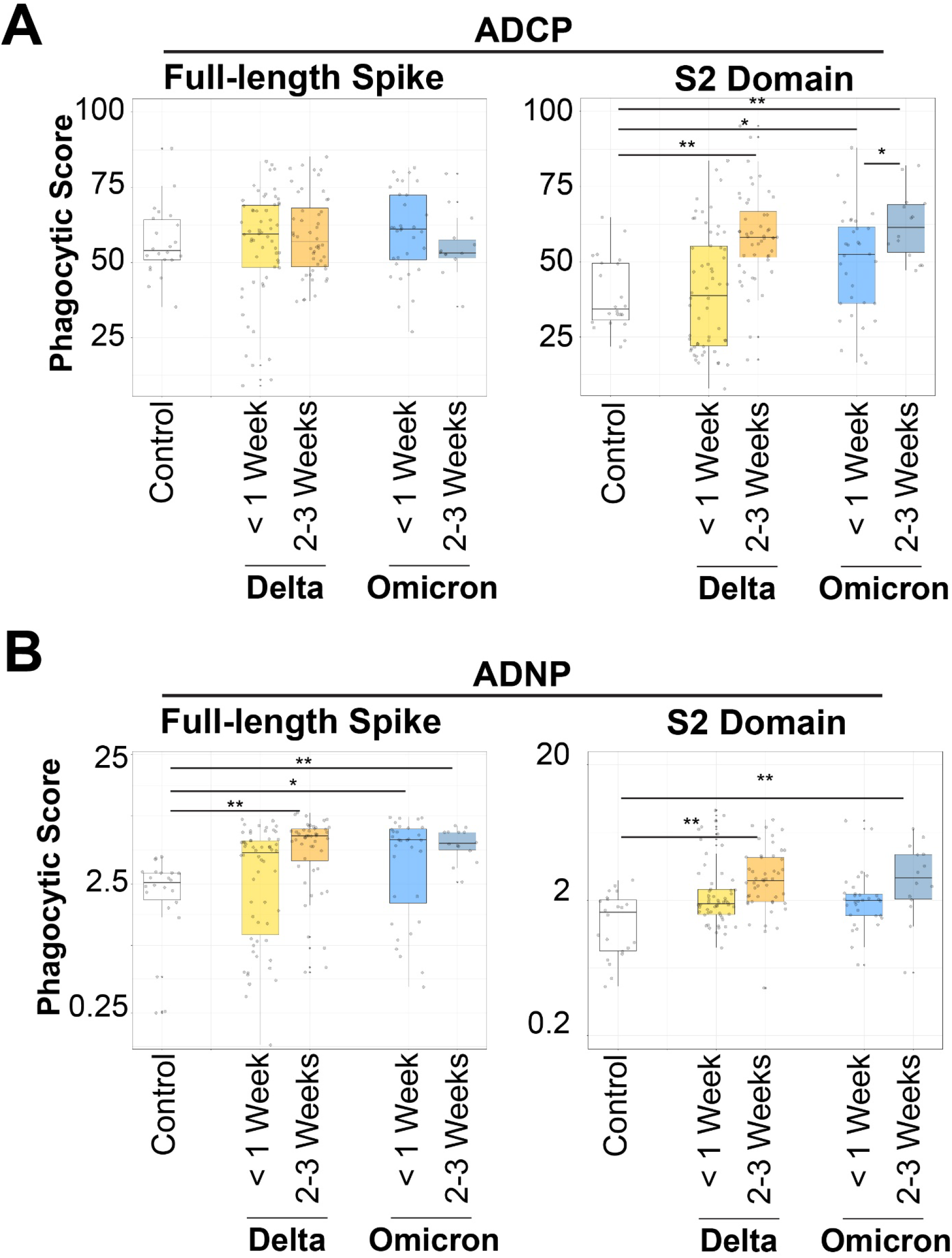
Expansion of S2 recognition is functionally linked to antibody-mediated opsinophagocytic activity. (A) Antibody-mediated cellular phagocytosis (ADCP) by monocytes was quantified using sera from vaccinated, non-breakthrough controls (column 1, white), vaccinated Delta breakthroughs at < 1 Week (yellow, column 2) or 2-3 Weeks post-breakthrough (orange, column 3), and vaccinated Omicron breakthroughs at < 1 Week (blue, column 4) or 2-3 Weeks post breakthrough (dark blue, column 5) for full-length Spike (left) and for the S2 domain alone (right). (B) Same as A, but for antibody-dependent neutrophil phagocytosis (ADNP) isolated from healthy donors. * = p < 0.05, and ** = p < 0.01 for all panels.

Antibody-mediated neutrophil phagocytosis (ADNP) has been linked to natural resolution of infection, convalescent plasma therapeutic activity, and vaccine-mediated immunity (*19, 24*). Interestingly, Spike-specific ADNP activity was not expanded over the first week of breakthrough infection in Delta breakthroughs, but did expand highly significantly over the following 2-3 weeks following infection. However, Spike-specific ADNP expanded more rapidly after breakthrough Omicron infection. Interestingly, S2-specific ADNP was not observed in either Delta or Omicron cases 1 week after breakthrough, but S2-specific ADNP responses expanded significantly over the next 2 weeks following infection (**Figure 4B**). Neutralization expansion of Spike increased but was correlated exclusively with the RBD domain (*25*). That neutralization did not correlate with S2, even though titers were bolstered, further highlights the non-neutralizing activity of S2-specific IgG1 post-breakthrough (Supplementary Figure 4). Thus, these data point to a simultaneous expansion of both neutralizing and opsinophagocytic antibodies, with a slightly earlier expansion of opsinophagocytic responses that may be key to the early capture, blockade, and elimination of the virus upon transmission.

### S2-expansion differences across mRNA platforms

Emerging data point to differences in Pfizer/BNT162b2 and Moderna/mRNA-1273 vaccine real world efficacy (*26*) and in immune Fc-profiles likely attributable to differences in dose, vaccine intervals, lipid-nanoparticle differences, and potential differences in mRNA chemistry (*27*). To determine if the 2 mRNA platforms induced a similar anamnestic response, breakthrough cases were split across individuals that received either of the mRNA vaccine platforms (**Figure 5**). Recipients of the Pfizer/BNT162b2 vaccine experienced a more rapid rise in Spike-specific IgG titers following breakthrough infection, although Pfizer/BNT162b2 and mRNA-1273 vaccinees reached similar Spike-specific titers 2-3 weeks after breakthrough infection (**Figure 5A**). Conversely, Pfizer/BNT162b2 vaccinees experienced a more significant and sustained increase in S2-specific immunity following breakthrough infection compared to Moderna/mRNA-1273 recipients (**Figure 5B**). However, despite this differential quantitative expansion, enhanced titer-corrected opsinophagocytic activity was observed following mRNA-1273 vaccination (**Figure 5C**-**D**), pointing to a distinct shift in responses across the 2 vaccine platforms, with a robust quantitative increase in S2-titers following Pfizer/BNT162b2 but more functional per-S2-specific antibody response following Moderna/mRNA-1273 vaccination.

**Figure 5.**
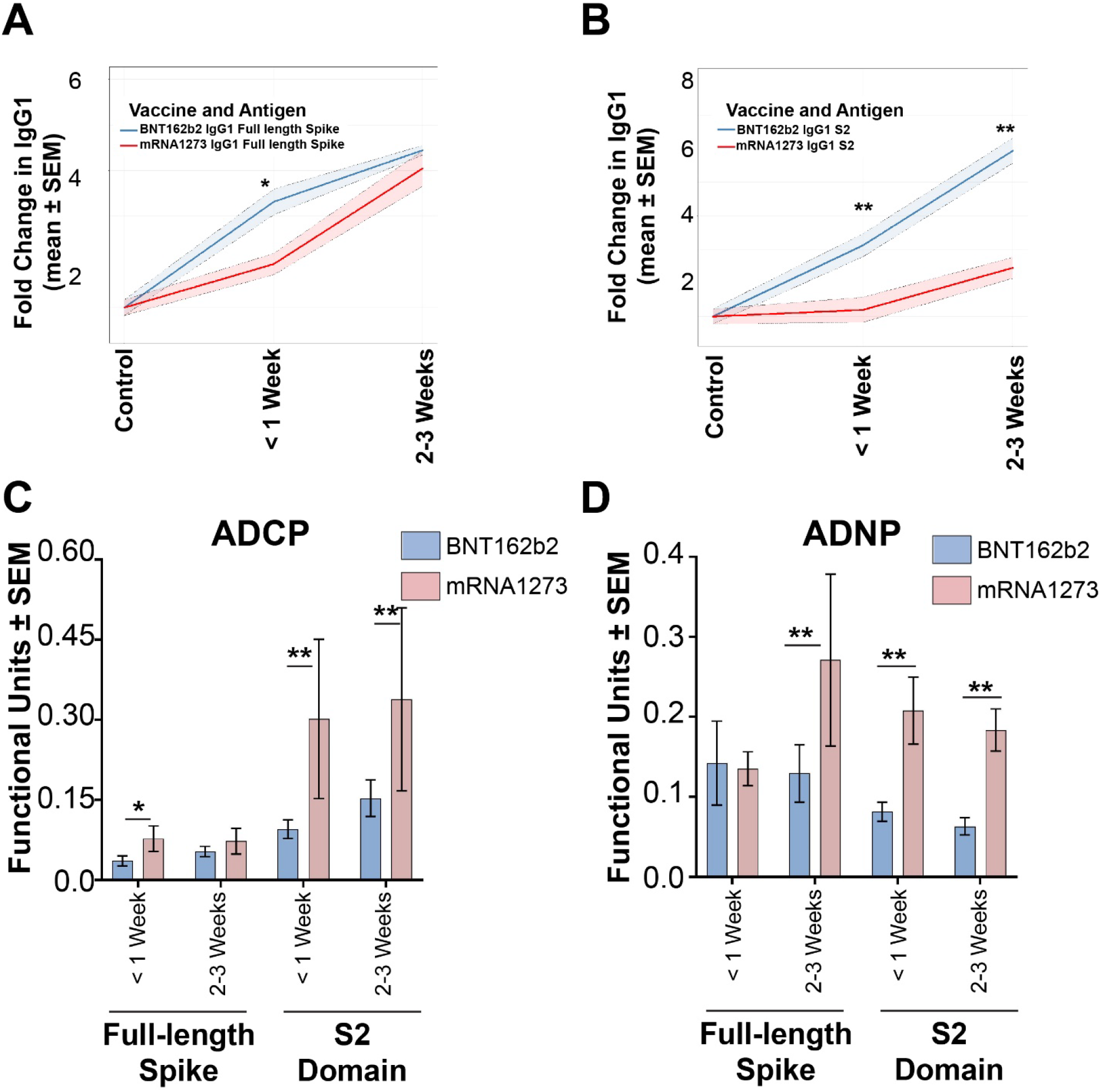
Breakthrough COVID-19 in mRNA vaccinated individuals results in distinct S2 functional expansions. (A) Fold changes in full-length Spike IgG1 in breakthrough COVID-19 cases in individuals vaccinated with the BNT162b2 (blue) or the mRNA1273 (red) mRNA vaccines at <1 Week or 2-3 Weeks post-breakthrough. (B) Fold changes in S2-specific IgG1 in breakthrough COVID-19 cases in BNT162b2 (blue) or the mRNA1273 (red) mRNA vaccine recipients at <1 Week or 2-3 Weeks post-breakthrough. ADCP functional units for individuals vaccinated with BNT162b2 (blue) or the mRNA1273 (red) mRNA vaccine against full-length Spike or S2. Functional units were quantified by taking the endocytic score divided by the MFI of IgG1 towards the antigen. Same as C, but for ADN* = p < 0.05, and ** = p < 0.01 for all panels.

### Selective expansion of fusion peptide and heptad repeat 1 specific immunity

To finally define whether S2-specific responses targeted particular regions of S2, we finally mapped the expanding breakthrough response across peptides spanning the S2 domain of the Spike antigen. Using peptides spanning regions across the S2-segment of the SARS-CoV-2 Spike antigen, a highly focused expansion of antibodies was noted to the second fusion peptide (FP2) and the heptad repeat 1 (HR1) (**Figure 6A**) across both Delta and Omicron breakthrough cases 7 days after breakthrough. While the response evolved to target additional regions of the S2 antigen in both groups over the first month following breakthrough infection, the FP2- and HR1-specific responses remained most highly immunodominant. At a more granular level, FP2 and HR1 IgG titers were already significantly higher than observed in controls after the first week of infection across VoC breakthrough groups (**Figure 6B-C**). Moreover, these responses remained persistently higher over the following 2-3 weeks. Lastly, we probed whether responses to particular subregions within S2 were correlated with virus clearance. Comparison of SARS-CoV-2 RNA decay slopes with S2-peptide-specific IgG expansions over the course of the first 3 weeks of breakthrough infection pointed to a highly significant inverse correlation between the slope of the evolution of FP2-specific responses and viral clearance (**Figure 6D**). Additionally, S2-peptide-specific responses were also significantly associated with viral clearance, pointing to a potentially critical role of multiple S2-specific responses in the elimination of viral replication. Collectively, these data point to a common immunodominant S2 anamnestic response to FP2, and the heptad repeats that may form the basis of a rapid functional humoral response required to capture, contain, and eliminate VoCs until T cells are able to traffic, expand, and eliminate remaining virus to ultimately clear the disease.

**Figure 6.**
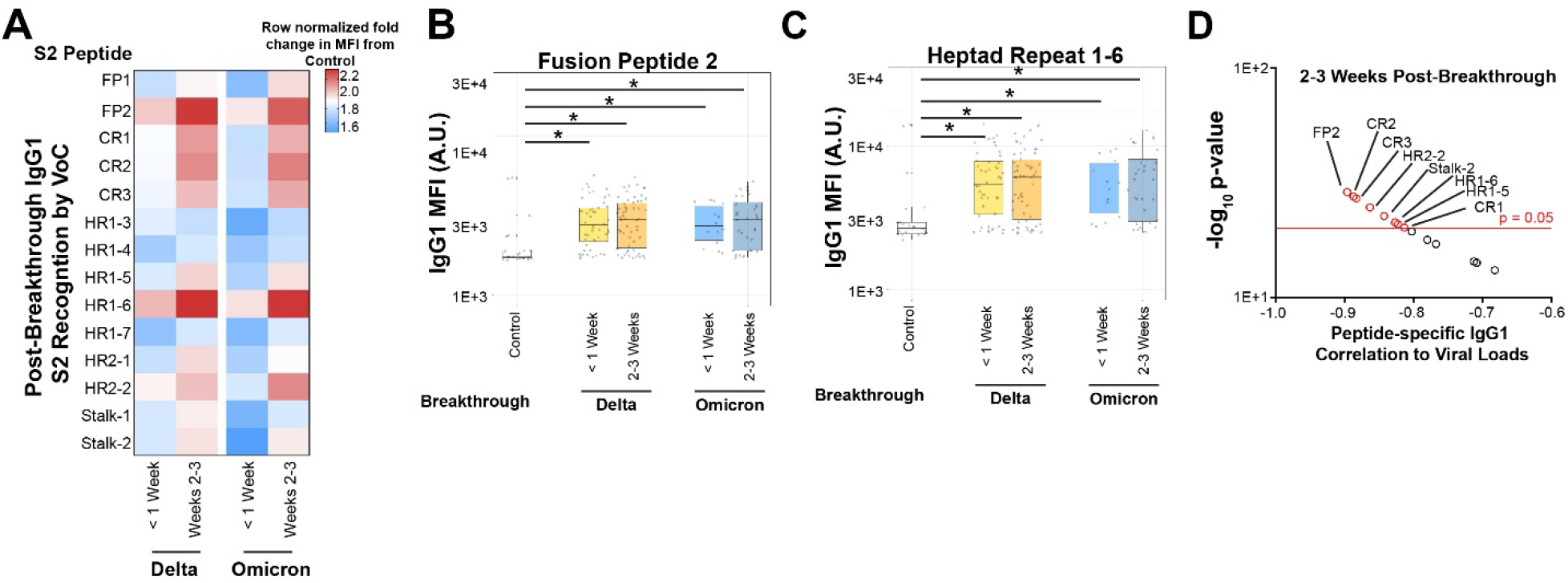
Immunodominant expansion of S2 is focused to the fusion peptide and heptad repeat region 1 for VoC breakthroughs. (A) Peptides spanning subregions of S2 were assayed for IgG1 expansion post-breakthrough by Delta (left two columns) and Omicron (right two columns) during the <1 Week and 2-3 Weeks post-breakthrough responses; heatmap legend is shown on the right. (B) The fusion peptide 2 (FP2) was assayed for IgG1 recognition in non-breakthrough, vaccinated controls (white, column 1), vaccinated Delta <1 Week (yellow, column 2) or 2-3 Weeks post-breakthrough (orange, column 3), and vaccinated Omicron <1 week (blue, column 4) or 2-3 Weeks post-breakthrough (dark blue, column 5). (C) Same as B, but for heptad repeat 1 subregion 6 (HR1-6). (D) Correlations between viral loads and IgG1 S2 peptide recognition for 2-3 Weeks post-breakthrough samples. * = p < 0.05.

## Discussion

Correlates of immunity represent immunological biomarkers that are statistically enriched in individuals that exhibit protective immunity in vaccine trials (*19, 28-32*). Correlates may be mechanistically involved in protective immunity, but can also represent surrogates of other immunological mechanisms key to anti-pathogen control. Most correlates of immunity are defined at the time of peak immunogenicity, aimed at defining immune responses able to predict clinical outcomes. However, within most trials, infections occur over prolonged periods of time, and vaccine-induced immune responses may have waned from peak immunogenicity. Thus, the immunological markers associated with protection over time may differ from those observed at peak immunogenicity. However, the collection of samples prior to infection over time in large Phase 3 trials is cost-prohibitive. Instead, the analysis of responses that selectively expand soon after infection can offer additional insights into the mechanism(s) by which vaccine-induced immunity reacts to control a challenge.

Phase 3 peak immunogenicity correlates analyses pointed to a strong association between vaccine-induced Spike-specific neutralizing and binding antibody titers with protection across the mRNA platforms (*6, 8*). Here we profiled the post-infection immunological profiles that evolved in breakthrough infections, with a unique interest in defining whether the kinetics of breakthrough correlates of immunity were consistent across variants of concern. While an expansion of full-length Spike IgG responses was observed in both Delta and Omicron breakthrough infections, limited expansion was observed in NTD-specific titers across both groups. Instead, the majority of the Spike-specific expansion was related to a unique anamnestic expansion of early S2 FP2- and HR1-specific IgM antibodies able to leverage monocyte phagocytosis, followed by a more mature S2 FP2- and HR1-specific IgG FcR binding neutrophil recruiting response observed both in Delta and Omicron breakthroughs. Thus, despite the immunodominant vaccine-induced response to the RBD, these data point to a critical and unexpected role of S2-specific functional humoral immunity as critical anamnestic correlates of immunity across VoCs.

Early studies of immune correlates of natural resolution of COVID-19 pointed to a selective expansion of S2-specific functional humoral immunity in survivors of natural severe COVID-19 infection. S2-specific antibodies were noted in survivors of severe COVID-19 from the time of intensive care unique admission (*22, 31*). Moreover, expanded S2-specific humoral immune responses were also noted in children (*32*), individuals that developed milder forms of COVID-19, as well as in individuals that developed asymptomatic infections (*31*). Interestingly, these natural S2-specific humoral immune responses may have emerged from pre-existing common-coronavirus specific humoral immune responses, that also expanded, marked by preferential FcR-binding, among individuals with asymptomatic infection. Common β-coronaviruses are largely conserved in their S2 domains. S2-specific monoclonals exhibit cross-coronavirus reactivity and *in vivo* protection in an Fc-dependent manner (*33*), arguing that these less potent neutralizing antibodies target a highly conserved region of the SARS-CoV-2 Spike may depend on non-neutralizing mechanisms of action. Given the robust association between pre-existing common-coronavirus immunity and attenuated COVID-19 (*32*), these data point to a critical role for cross-reactive S2-specific humoral immunity in both infection and vaccine-induced immune responses.

The S2 includes the fusion machinery required for viral entry (*34*), requiring precise packaging and movement upon Spike binding to the host angiotensin-2 (ACE2) receptor. Thus, unlike other regions of the Spike antigen, the S2 is less mutable and has shown conservation across distinct VOCs, with only ∼2 and 12 mutations in Delta and Omicron respectively (depending on sublineage), representing the most highly conserved region of the SARS-CoV-2 Spike antigen. However, the S2-directed response is less dominant following mRNA vaccination, likely related to the 2 proline stabilization introduced in the protein to stabilize the antigen during vaccination (*35, 36*). This stabilization holds S1 and S2 in a pre-fusion state, likely required to drive robust immunity to the primary target of neutralizing antibodies, the RBD. However, this stabilization also may make S2 less accessible to the immune response. Infection with the virus generates copious amounts of Spike in its inherently destabilized form. This allows immune surveillance networks to sample both S1 and S2 domains in pre- and post-fusion forms, triggering an anamnestic response. The differences in the anamnestic immunodominance of S1 and S2 may relate to the fact that S1 is presented largely as soluble protein, whereas S2 may be presented in a particulate form, due to the C-terminal transmembrane anchoring domain of the protein. Conversely, S2-responses may gain a competitive advantage due to the high degree of conservation across VoCs, able to recruit pre-existing B cells, whereas previously programmed RBD-and NTD-specific B cells may struggle to bind to the incoming VoC due to significant antigenic variation (*1, 2, 4*). However, despite the enhanced sequence conservation between the vaccine strain and Delta compared to Omicron, S2-specific immunity expanded in both Delta and Omicron breakthroughs, suggesting that S2-specific anamnestic correlates may be key to protection across VoCs. Differences in dose, vaccination interval, lipid-nanoparticle composition, and mRNA chemistry all contribute to differences in antibody subclass/isotype and Fc-receptor binding profiles across mRNA vaccines (*27*). Here we observed differences in the anamnestic response following Pfizer/BNT162b2 and Moderna/mRNA-1273 immunization, linked to a higher magnitude expansion of anamnestic immunity in Pfizer/BNT162b2 vaccinees and a functional expansion in Moderna/mRNA1273 vaccinees, although both breakthrough profiles resulted in an expansion of S2-specific immunity. These differences may be related to real-world efficacy differences across the platforms, with reduced breakthrough infections observed in Moderna/mRNA1273 vaccinees, potentially related to higher IgA titers and functional humoral immunity (REF) that may provide a more robust barrier against infection at the mucosal barrier. However, the expansion of S2-specific immunity following both vaccines, focused on FP2 and HR1, may be related to their accessible positions on the Spike antigen (*37-39*). Thus, strategies to boost functional humoral immunity to these critical sights may represent a promising future strategy to maintain long-term protection against severe disease and death against current and future VoCs. Whether these S2-specific responses can work in concert with T cells that also target conserved regions of the SARS-CoV-2 Spike remains unclear (*12*); however, these responses point to important, unexpected targets of the immune response that may be key to the durable protection against disease severity. Moreover, these regions that show higher conservation between VoCs, and β-coronaviruses in general, that are expanded post-infection could be viewed as rational vaccination booster targets.

## Methods

### Study Approval

Approval for study for breakthrough COVID-19 at Massachusetts General Brigham was approved under protocol 2021P000812 by the Mass General Brigham IRB. For this cohort, symptomatic COVID-19 patients seen for outpatient care were recruited based on a positive SARS-CoV-2 test (**Table S1**). Sequencing from anterior nasal swabs was performed for VoC identification. The use of healthy donor blood for cellular functional assays is approved under protocol 2021P002628 by the Mass General Institutional IRB.

The Hospitalized or Ambulatory Adults with Respiratory Viral Infections (HAARVI) study was approved by the University of Washington Human Subjects Division Institutional Review Board (STUDY00000959).

### Antigens

All antigens and peptides used in this study are listed in **Table S2** and **Table S3**. The protein antigens were received in lyophilized powder form and resuspended in water to a final concentration of 0.5 mg/mL. Peptides were received in solution and, if necessary, were buffer exchanged using Zeba-Spin columns (ThermoFisher, USA).

### Immunoglobulin isotype and Fc receptor binding

Sera was collected from participants at two time points post-COVID-19 diagnosis, with the first time point being 6.3 ± 2.8 days post-observed start date, and 18.8 ± 2.9 days post-observed start date. The two groups were clustered together for analyses and classified as < 1 Week and 2-3 Weeks.

Systems serology for antigen-specific recognition was done using custom multiplex magnetic Luminex beads (Luminex Corp, TX, USA) as previously as previously described (*21*). Antigens and peptides were coupled to beads through carbodiimide-NHS ester-coupling chemistry. The antigen-or peptide-coupled beads were incubated with heat-inactivated serum (1:100 for IgG2, IgG3, IgG4, IgM, and IgA1, 1:250 for IgG1, and 1:750 for Fcγ-receptor binding) overnight at 4°C in 384 well plates (Greiner Bio-One, Germany). Secondary antibodies were PE-conjugated and incubated with samples at room temperature for 1 hour at a 1:100 dilution in sterile-filtered Assay Buffer (1X PBS, pH = 7.4, 0.1 % BSA, 0.05 % Tween – 20). For Fcγ-receptors, PE-streptavidin (Agilent Technologies, CA, USA) at a 1:1000 dilution.

For flow cytometry analysis, the IQue Screener PLUS cytometer (IntelliCyt) was used using customized gating for each bead region. Fluorescence in the BL2 channel was quantified and exported into .csv files and subsequently analyzed (see below).

### Viral loads quantitation

SARS-CoV-2 RNA was extracted and quantified as previously described (*20, 40*). VoC identification through sequencing was done using a previously validated method (*25*). Peak viral loads for each patient were quantified and the mean peak viral loads between VoC was calculated.

### Evaluation of antibody-mediated functions

Antibody-dependent cellular phagocytosis (ADCP) by monocytes and neutrophil phagocytosis (ADNP) were quantified using a validated, flow cytometry-based bead phagocytic assay. Fluorescently labeled microspheres were coupled to antigens through biotinylation and conjugation to neutravidin beads. Diluted and heat-inactivated serum samples were incubated with the antigen-coupled neutravidin beads to create a pre-immune complex. The solution was then incubated with THP-1 monocytes (ATCC, Manassas, USA) or primary-derived neutrophils (*21*). For ADNP, cells were stained with anti-CD66b Pac blue antibody to calculate the percentage of CD66b+ neutrophils. Cells were fixed with 4% paraformaldehyde. Microsphere uptake was quantified by the percentage of microsphere-positive cells × MFI of microsphere-positive cells.

### Statistical Analyses

Data visualizations and analysis were done using R Studio V 1.4.1103 or GraphPad Prism. Box and whisker plots were generated using ggplot showing the mean and standard deviation for each group as factors. An initial ANOVA was performed to identify significant groupings. A Wilcoxon-rank sum test and pairwise T-tests were used to determine grouping between two groups. For all analyses * stands for p < 0.05, and ** stands for p < 0.01. All codes and scripts are available upon request and no original code was created for this manuscript.

## Acknowledgments

We thank Mark and Lisa Schwartz, Terry and Susan Ragon, and the SAMANA Kay MGH Research Scholars award for their support. GA also receives funding from the Massachusetts Consortium on Pathogen Readiness (MassCPR), the Gates Global Health Vaccine Accelerator Platform, and the NIH (3R37AI080289-11S1, R01AI146785, U19AI42790-01, U19AI135995-02, U19AI42790-01, P01AI1650721, U01CA260476 – 01, CIVIC75N93019C00052).

## Disclosure

Galit Alter is a founder/equity holder in Seroymx Systems and Leyden Labs. GA has served as a scientific advisor for Sanofi Vaccines. GA has collaborative agreements with GSK, Merck, Abbvie, Sanofi, Medicago, BioNtech, Moderna, BMS, Novavax, SK Biosciences, Gilead, and Sanaria.

