## Supplemental Information for "Anamnestic Humoral Correlates of Immunity Across SARS-CoV-2 Variants of Concern"

Includes Supplementary Figures 1-4, Supplementary Figure Legends, and Supplementary Tables 1-3.

**Supplementary Figure 1**

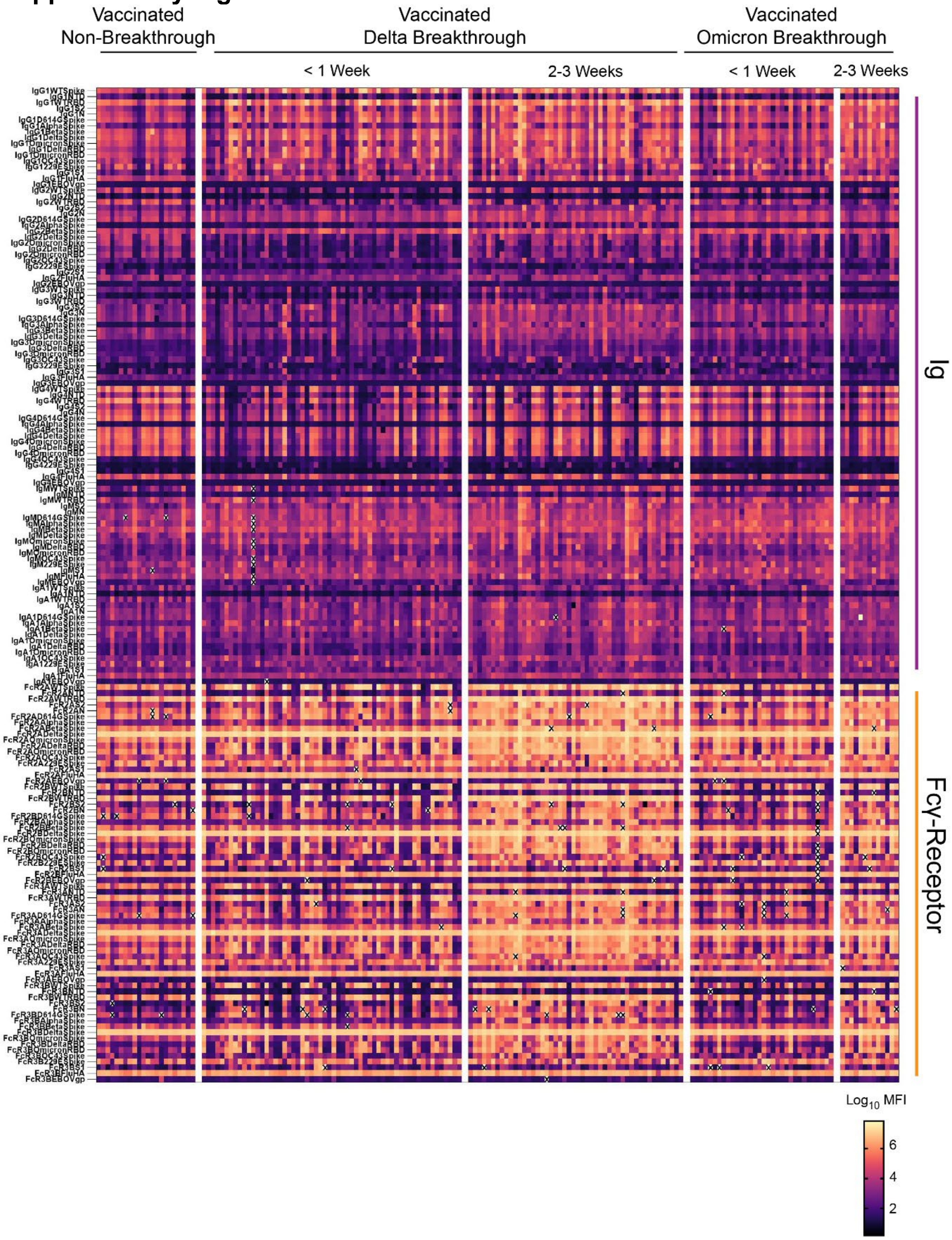


**Supplementary Figure 1. Biophysical systems serology of antibody and Fcγ-receptor binding to antigens for breakthrough COVID-19.** Sera from vaccinated, non-breakthrough controls, Delta breakthroughs, and Omicron breakthroughs were analyzed by systems serology for recognition of various antigens during < 1 Week and 2-3 Weeks post-observation following breakthrough. Each column represents a single patient within the specified cohort. Gaps are shown to distinguish groupings on the top, and side panels are shown to distinguish between Ig and FcR binding. Shown on the bottom right is the heatmap legend.

**Supplementary Figure 2**

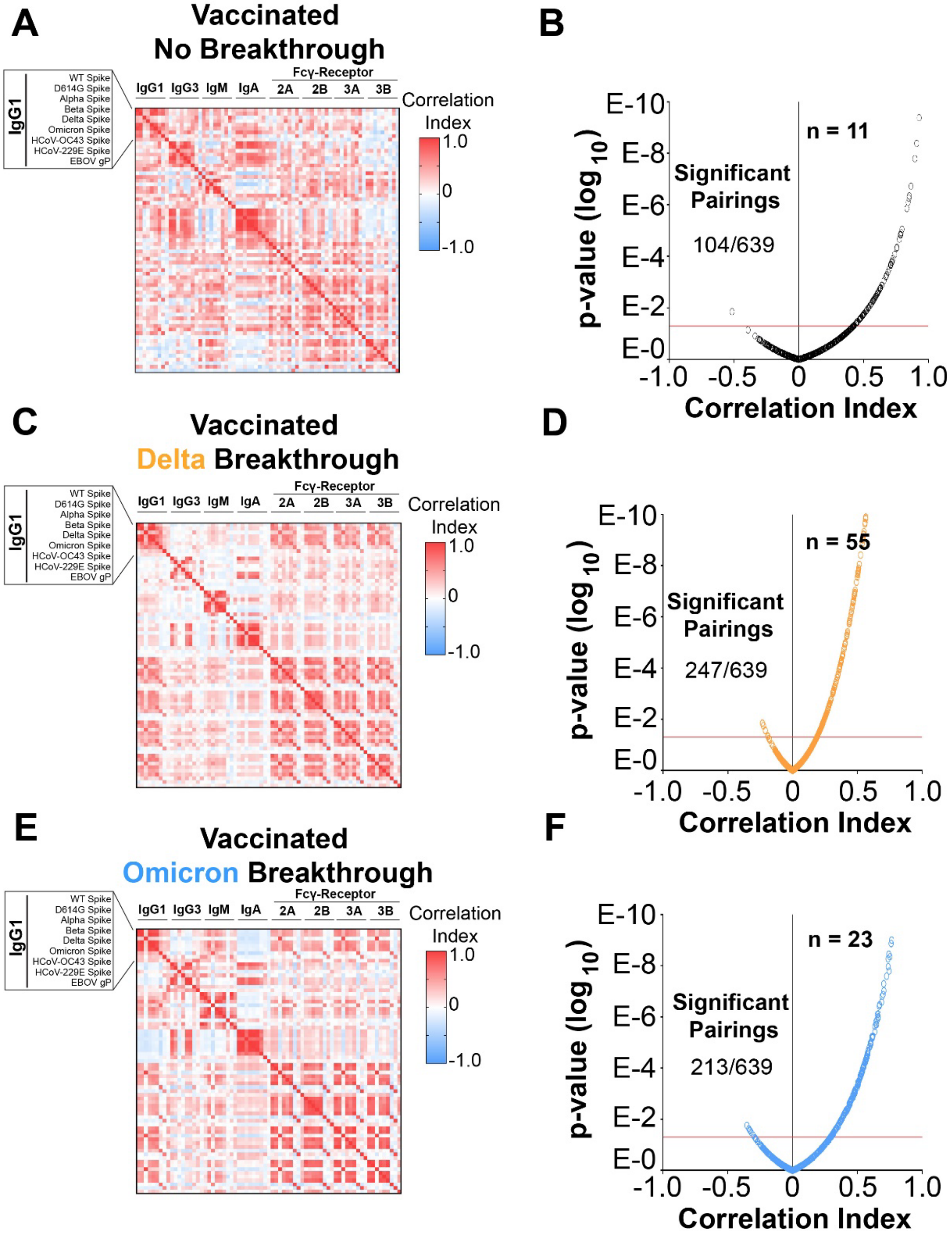


**Supplementary Figure 2. Breakthroughs in vaccinated individuals expanded correlates of VoC recognition.** (A) Correlations between antibody and Fc-effector recognition of SARS-CoV-2 VoCs in vaccinated, non-breakthrough controls. Shown in the upper left is a zoom-in view of the ordering of antigens, which is repeated for IgG1, IgG3, IgM, IgA1, FcγR2A, FcγR2B, FcγR3A, and FcγR3B. Ebola virus glycoprotein (EBOV gP) is used as a negative correlate control. Shown on the right is the correlation heatmap legend. (B) Volcano plot of pairwise correlations for vaccinated, non-breakthrough controls. The red line on the y-axis represents statistical significance cutoff (p = 0.05). Shown in the graph are the number of statistically significant correlates. (C) Same as A, but for vaccinated Delta breakthroughs 2-3 Weeks post-breakthrough. (D) Same as B, but for vaccinated Delta breakthroughs 2-3 Weeks post-breakthrough. (E) Same as A, but for vaccinated Omicron breakthroughs 2-3 Weeks post-breakthrough. (F) Same as B, but for vaccinated Omicron breakthroughs 2-3 Weeks post-breakthrough.

### Supplementary Figure 3

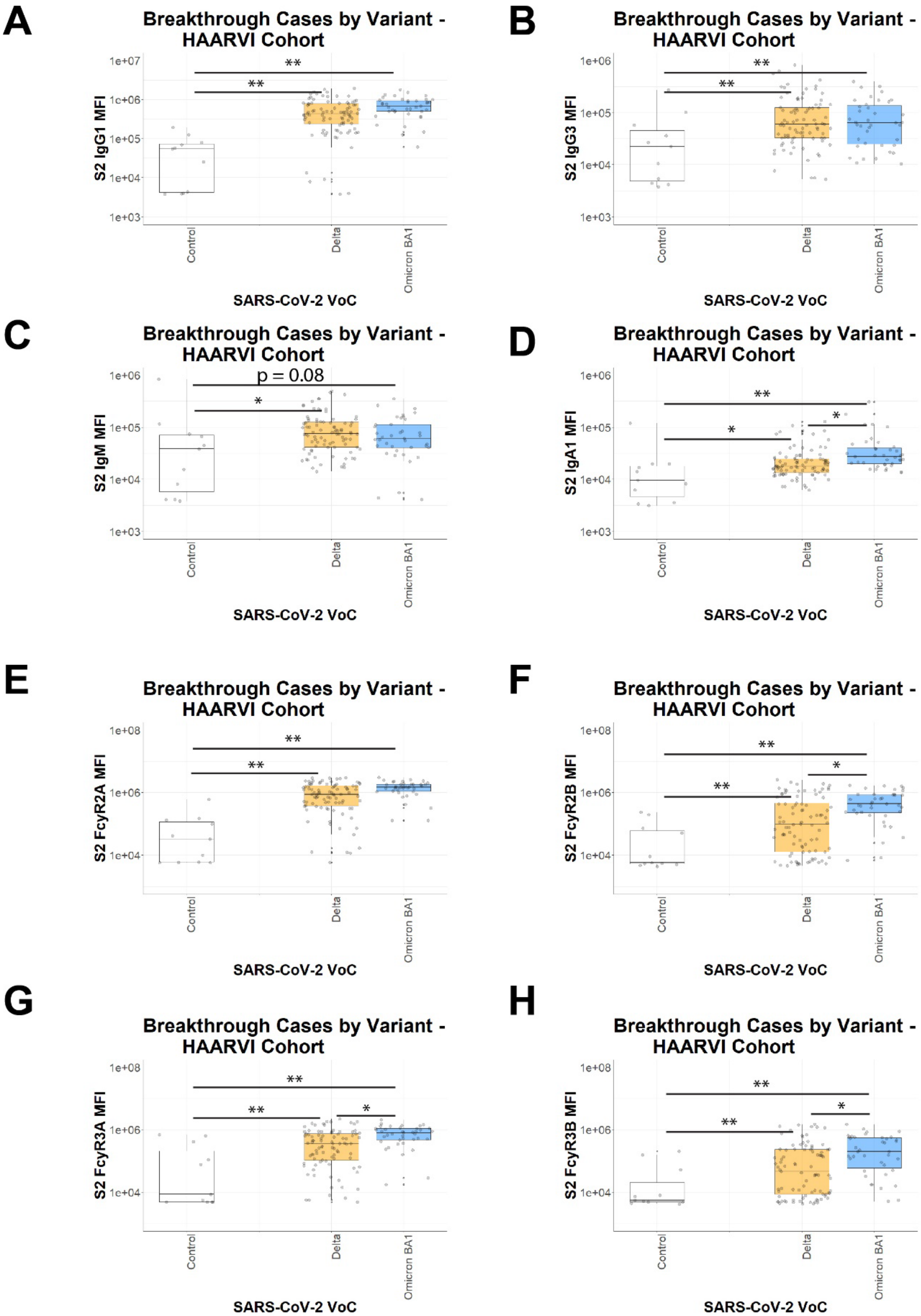

**Supplementary Figure 3. S2-specific antibodies are expanded in breakthrough infections in an independent cohort.** (A) Uninfected controls (column 1, white), Delta breakthroughs (column 2, orange), and Omicron BA.1 (blue, column 3) were assayed for S2-specific IgG1. Sera were harvested 14 days after diagnosis. (B) Same as A, but for IgG3. (C) Same as A, but for IgM. (D) Same as A, but for IgA1. (E) Same as A, for S2-specific FcγR2A responses. (F) Same as E, but for FcγR2B. (G) Same as E, but for FcγR3A**.** (H) Same as E, but for FcγR3B. * = $p < 0.05$, and ** = $p < 0.01$ for all panels.

**Supplementary Figure 4**

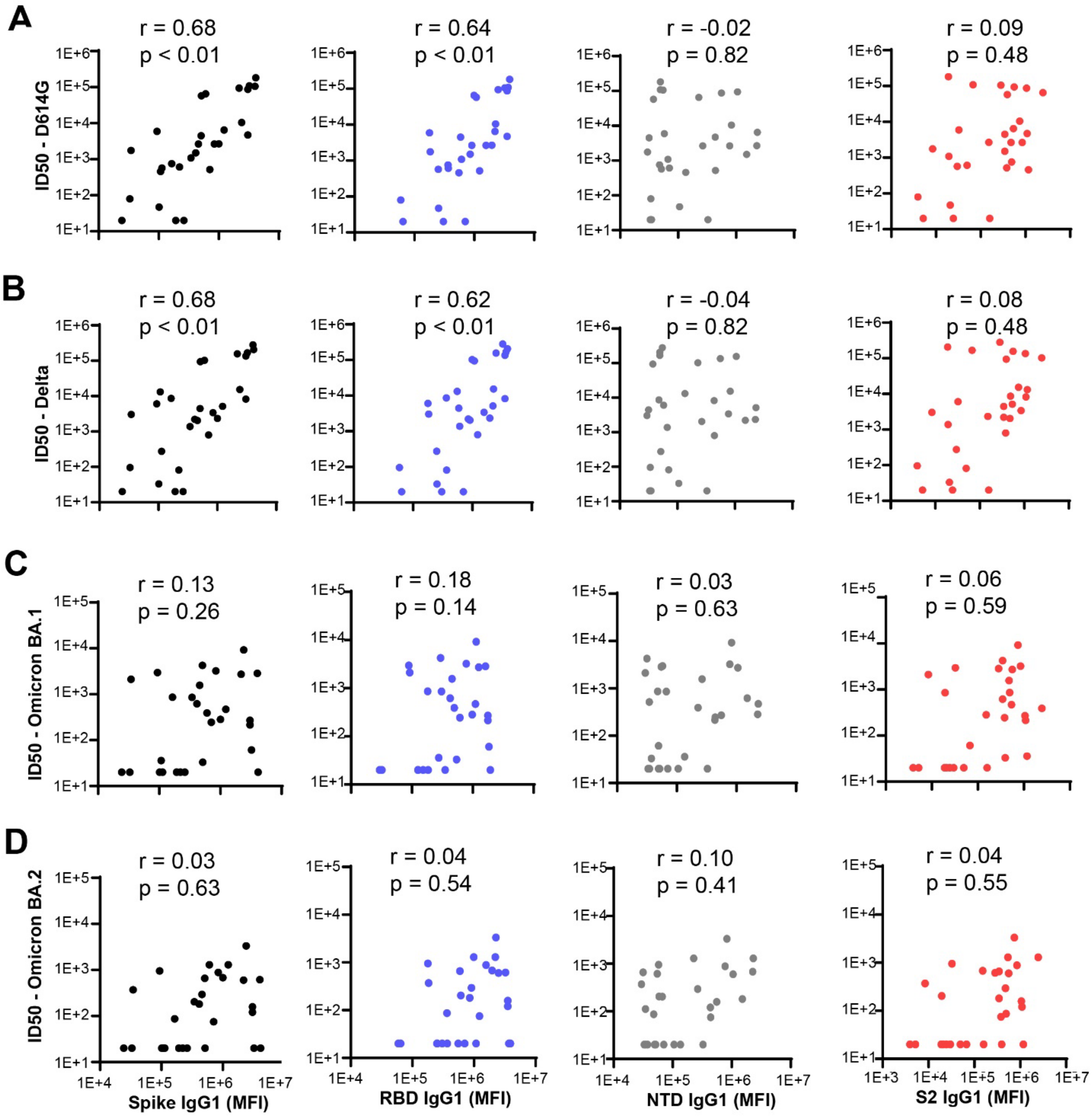


**Supplementary Figure 4. Correlations between neutralization and Spike domains are focused to the RBD of D614G and Delta, but not to S2, post-breakthrough**. (A) Vaccinated individuals with breakthrough COVID-19 were assayed for correlation between neutralization (ID50) and IgG1 Spike (black), RBD (blue), NTD (gray), or S2 (red). Shown above each panel are correlation coefficients and p-values. (B) Same as A, but for neutralization of Delta. (C) Same as A, but for neutralization of Omicron BA.1. (D) Same as A, but for neutralization of Omicron BA.2

**Table S1**

| | Control (Vaccinated, non-breakthrough) | Vaccinated, Delta VoC Breakthrough | Vaccinated, Omicron VoC Breakthrough |
|---|---|---|---|
| N | 11 | 37 | 19 |
| Peak Viral Loads (log10) | N/A/ | 6.2 ± 1.5 | 5.9 ± 1.6 |
| Fully Vaccinated | 11 | 28 | 18 |
| Breakthrough from mRNA-1273 | N/A | 6 | 6 |
| Breakthrough from BNT162b2 | N/A | 14 | 9 |

**Table S1. Cohort analysis by VoC breakthrough and mRNA vaccine type.**

**Table S2**

| Item | Supplier | Catalog Number |
|---|---|---|
| Anti-Human IgG1-PE | Southern Biotech | HP6001 |
| Anti-human IgG2-PE | Southern Biotech | 31-7-4 |
| Anti-human IgG3-PE | Southern Biotech | HP6050 |
| Anti-human IgG4-PE | Southern Biotech | HP6025 |
| Anti-human IgM-PE | Southern Biotech | SA-DA4 |
| Anti-human IgA1-PE | Southern Biotech | HP6025 |
| SARS-CoV-2 WT Spike | Sino Biological | 40589-V08H4 |
| SARS-CoV-2 WT S1 Domain | Sino Biological | 40591-V08H |
| SARS-CoV-2 WT Receptor Binding Domain (RBD) | Sino Biological | 40592-V08H |
| SARS-CoV-2 WT S2 Domain | Sino Biological | 40590-V08B |
| SARS-CoV-2 WT N-terminal Domain | Sino Biological | 40591-V49H |
| SARS-CoV-2 Alpha Variant S | Sino Biological | 40589-V08B6 |
| SARS-CoV-2 Beta Variant S | Sino Biological | 40589-V08B7 |
| SARS-CoV-2 Gamma Variant S | Sino Biological | 40589-V08B10 |
| SARS-CoV-2 Delta Variant S | Sino Biological | 40589-V08B16 |
| SARS-CoV-2 Omicron Variant S | Sino Biological | 40589-V08H26 |
| Human Coronavirus OC43 S | Sino Biological | 40607-V08B |
| Human Coronaivirus HKU1 S (isolate N5) | Sino Biological | 40606-V08B |
| Human Coronavirus 229E S | Sino Biological | 40605-V08B |
| Human Cytomegalovirus (HCMV) Glycoprotein B (gB) | Sino Biological | 10202-V08H1 |
| Ebola Virus Glyoprotein | IBT Bioservices | 0501-015 |
| Anti-Human IgG1-PE | Southern Biotech | HP6001 |
| Anti-human IgG2-PE | Southern Biotech | 31-7-4 |
| Anti-human IgG3-PE | Southern Biotech | HP6050 |
| Anti-human IgG4-PE | Southern Biotech | HP6025 |
| Anti-human IgM-PE | Southern Biotech | SA-DA4 |
| Anti-human IgA1-PE | Southern Biotech | HP6025 |
| Ebola Virus Glyoprotein | IBT Bioservices | 0501-015 |
| PE-Streptavidin | Agilent Technologies | PB32-10 |
| NHS-Sulfo-LC-LC Kit | ThermoFisher | 21435 |
| Zebra-Spin Desalting and Chromatography Columns | ThermoFisher | 89882 |
| MagPlex Microspheres | Luminex MFG | MC12001-01 (Cataloged by region) |
| R Studio V 1.4.1103 | RStudio, PBC | Open Source |
| GraphPad Prism | GraphPad Software, LLC | Ragon Site License |
| FlowJo V. 10.8 | FlowJo, LLC | www.flowjo.com/solutions/flowjo/downloads |
| iQue Forecyt | Sartorius | 60028 |
| iQue Screener Plus | Intellicyt/Sartorius | 11811 |

| | | |
|---|---|---|
| 384-well HydroSpeed Plate Washer | Tecan | 30190112 |

**Table S2. Table of the reagents, analysis software, and instrumentation used in this study.**

**Table S3**

| Peptide Name | Sequence |
|---|---|
| FP1 | DPSKPSKRSFIEDLLFNKV |
| FP2 | SFIEDLLFNKVTLADA |
| CR1 | QYGDCLGDIAARDLICAQKFNG |
| CR2 | LTDEMIAQYTSALLAGTITSGWTFGAGA |
| CR3 | IPFAMQMAYRFNGIG |
| HR1-3 | KLIANQFNSAIGKIQDSLSSTASALGKLQDVVNQN |
| HR1-4 | ALGKLQDVVNQNAQALNTLVKQLSSN |
| HR1-5 | AISSVLNDILSRLDKVEAEVQ |
| HR1-6 | QLSSNFGAISSVLNDILSRLDKVEAEVQIDRLITGRLQS |
| HR1-7 | SVLNDILSR |
| Stalk-1 | DPLQPELDSFKEELDKYFKNHTSPD |
| Stalk-2 | SFKEELDKYFKNHTS |
| HR2-1 | ASVVNIQKEIDRLNEVAKNLNESLIDLQELGKYEQ |
| HR2-2 | QKEIDRLNEVAKNLNESLIDLQE |

**Table S3. S2 Peptide names and sequences used for this study.**
